# An index-based system for early alerts of potential zoonotic disease outbreaks

**DOI:** 10.1101/2025.06.17.660271

**Authors:** Jaqueline S. Angelo, Livia Abdala, Douglas A. Augusto, Marcia Chame, Eduardo Krempser

## Abstract

The Information System on Wildlife Health (SISS-Geo) is a free platform designed for the real-time collection of georeferenced data on wildlife health and environmental conditions via mobile devices. It serves as a collaborative tool, enabling health professionals, researchers, environmental managers, and the general public to report information about wildlife health occurrences directly within the system. Since its launch in 2014, SISS-Geo has been successfully applied in supporting decision-making during significant wildlife health events in Brazil. In this paper, we introduce a dynamic alert system based on a Multi-Attribute Zoonotic Alert Index (Z-Alert) to enhance disease outbreak detection and response in wildlife. The proposed approach improves the current alert system of the SISS-Geo platform by integrating clustering techniques with multi-objective optimization. The clustering stage allows to group SISS-Geo records in a way that reflects both spatial and temporal variations, capturing relevant patterns for epidemiological surveillance. Each identified cluster is assigned a numerical alert index, the Z-Alert index, reflecting its criticality level, based on optimally weighted attributes such as the percentage of dead animals, temporal interval, and geographical spread. The system offers customization alert thresholds, allowing health managers to adapt the alert system to their specific needs. To validate the model, we used historical yellow fever case data from Brazil’s Ministry of Health, achieving over 94% accuracy in identifying critical clusters. By enabling the prioritization of prevention and investigation actions, the system strengthens the responsiveness of health managers, optimizing resource allocation and promoting smart economic strategies for outbreak containment.

## Introduction

It is widely recognized that most emerging zoonotic diseases—those naturally transmissible from animals to humans— originate in wildlife and are driven by human activities like land-use changes, urbanization, and globalization. A recent review highlights that approximately 75% of all emerging infectious diseases are of zoonotic origin [22]. The effective surveillance and disease prevention requires a collaborative, multidisciplinary approach to understand disease ecology and conduct risk assessments. In this context, wildlife monitoring becomes essential for various sectors of society, especially for public health. Collecting and analyzing data on wildlife populations can provide valuable insights to guide preventive actions and control zoonotic disease transmission. The One Health approach is particularly relevant in this context, as it recognizes the interconnection between human, animal, and environmental health [23, 11]. This integrated perspective is especially important in a country like Brazil, where environmental and epidemiological complexities require comprehensive strategies for disease surveillance systems [8].

The effective implementation of One Health principles requires innovative technologies that facilitate real-time, high-quality data collection and support decision-making through efficient information analysis and management. To address this need, the Brazilian Wildlife Health Information System, SISS-Geo (*Sistema de Informação em Saúde Silvestre Georreferenciado (in Portuguese)* [3, 2] was created in 2014 by the Institutional Platform for Biodiversity and Wildlife Health at Oswaldo Cruz Foundation (Fiocruz) to support collaborative monitoring.

The SISS-Geo (avaliable at https://www.biodiversidade.ciss.fiocruz.br or https://sissgeo.lncc.br) is a digital platform that enables the real-time collection of georeferenced observations on wildlife and environmental conditions via mobile devices, with active public participation. It integrates the registration of occurrences, expert diagnoses, and computational tools for data analysis, prediction, visualization, and real-time alerts for wildlife-related health events. Since its launch, SISS-Geo has proven to be a valuable resource for supporting decision-making during major wildlife health events in Brazil, such as Yellow Fever (YF) outbreaks [20, 1, 9], by accelerating information dissemination, providing accurate geographic locations, and simplifying response efforts.

The One Health Joint Plan of Action (2022–2026) highlights the SISS-Geo as an example of an integrated surveillance system that supports early detection and response to zoonotic threats through multisectoral collaboration [7]. This positions SISS-Geo in alignment with the global call to implement the One Health Early Warning and Response System (OH-EWRS) approach for addressing emerging zoonotic diseases [10].

In Brazil, the YF virus is endemic to the Amazon region, eventually reemerging into epidemic waves [1]. While currently confined to South and Central America and parts of Africa, its potential for geographic expansion, raises concerns for public health and cross-border transmission [12]. This underscores the urgent need to develop early warning and response systems capable of detecting outbreaks in their initial stages and enabling timely, coordinated interventions. By enabling the real-time collection of high-quality data, the platform provides critical information to government agencies, the health sector, and society at large—even in some of the most remote regions of the country—thereby strengthening surveillance and response efforts across multiple levels.

One of the most strategic components of SISS-Geo’s features is the alert system, which notifies health managers about reported cases. Currently, the system follows fixed monitoring rules. For instance, if a non-human primate (NHP) death or illness is reported, an email is automatically sent to health managers registered on the platform along with information about the record. While this enables managers to make timely decisions regarding the actions to be taken, it is unable to detect levels of risk, as fixed rules fail to account for regional variability and complexity. Without an appropriate prioritization mechanism, it becomes more challenging to direct efforts to the most impactful cases, compromising the efficiency of investigation and intervention measures.

The identification and prioritization of actions in critical health regions are essential for the effective allocation of resources in disease prevention and control efforts. Such an approach prevents the waste of supplies and personnel in low-risk areas, focusing efforts where there is the greatest need. Timely detection of diseases through surveillance, combined with effective implementation of targeted interventions, can significantly reduce the scale, severity, and economic impact of responses to outbreaks [25].

In this study, we aim to leverage data from the SISS-Geo to investigate the potential of animal-related information as an early warning signal for disease outbreaks that may later affect humans. Our focus is on NHPs, which play a key role as sentinels for YF circulation. However, the proposed methodology is not limited to the yellow fever context, as the SISS-Geo system also contains records related to other infectious diseases, such as rabies and avian influenza. By analyzing spatial and temporal patterns of NHP cases, we seek to identify regions that exhibit early signs of viral activity. Using NHPs as indicators of environmental health risks enables earlier detection and supports timely public health responses before human infections are reported.

As part of the efforts to improve surveillance strategies, we propose an index-based alert system that incorporates regional information to accurately reflect the criticality of events. By creating a Multi-Attribute Zoonotic Alert Index (named Z-Alert), we aim to identify not only individual records but also regions with potential risks to human and animal health. The Z-Alert proposal is part of a broader effort to strengthen national surveillance strategies, especially following the expansion of YF into extra-Amazonian regions starting in 2014, which highlighted the need to enhance wildlife health surveillance in Brazil. In this context, tools such as SISS-Geo and the environmental suitability models developed by the Institutional Platform for Biodiversity and Wildlife Health (Pibss), in partnership with the Yellow Fever Modeling Group (GruMFA), have become central elements for anticipating risk areas, guiding vaccination efforts, and supporting the definition of ecological transmission corridors. Both have been officially incorporated into national response strategies, including epidemiological bulletins, technical notes [16, 17, 18] issued by the General Coordination for Arbovirus Surveillance of the Department of Health Surveillance and Environment of the Ministry of Health (CGARB/SVSA/MS), and the Contingency Plan for Public Health Emergencies – Yellow Fever (2nd edition) [19, 14].

Unlike the current rule-based model, the proposed system generates alerts based on clustered data analysis, considering multiple observations recorded in SISS-Geo. By integrating attributes derived from the platform’s georeferenced records, such as the number and proportion of dead animals, event duration, and geographic extent, the system identifies areas of greater concern. We combine clustering techniques—to capture spatial and temporal disease occurrence patterns—with an index-based alert system that measures alert severity. Such an approach enhances the capacity for early detection and response within the context of zoonotic disease surveillance, representing an evolution of the current rule-based notification scheme of SISS-Geo. With the aid of a data clustering technique and a multi-objective optimization process, the zoonotic alert system will integrate multiple attributes to estimate the health risk level associated with an event. We also propose customized alert thresholds that dynamically adjust the number of alerts/notifications generated by the system, ensuring that only the most relevant situations are reported to decision-makers.

In this paper, we aim to:

‐ establish an index-based system that reflects a level of risk and provides a more refined assessment of potential health risks. The alert system is capable of distinguishing varying levels of risk by incorporating multiple regional attributes.
‐ allow health managers to customize the alert system, enabling them to prioritize alerts and optimize investigation and intervention efforts according to their specific needs and realities. In a vast country like Brazil, where environmental, epidemiological, and logistical conditions vary widely, such flexibility is essential for efficient resource allocation.
‐ utilize clustering methods to capture spatial and temporal patterns of disease occurrence, enabling a more comprehensive analysis of outbreak dynamics.
‐ apply a multi-objective optimization process to balance key criteria, ensuring that alerts are accurate and minimize false negatives to prevent the non-identification of critical events while reducing false positives to avoid overloading verification systems.
‐ finally, but not limited to, strengthen the integration of human, animal, and environmental health data to improve zoonotic disease surveillance and prevention.

The system’s effectiveness relies heavily on the accuracy of its prediction model in correctly identifying both true positives (alerts) and true negatives (non-alerts). Missing an alert condition (false negative) can have serious consequences for wildlife, the environment, and human health. Conversely, false positives could overwhelm the limited network of laboratories and experts responsible for verifying alerts. Thus, finding the right balance between classifying alerts and non-alerts is crucial to ensuring that the system effectively prioritizes critical regions without overloading resources. In addition, the customization of alert thresholds plays a key role in this trade-off, as stricter limits reduce false positives but increase the risk of missing real cases (less sensitivity), while more relaxed limits enhance detection at the cost of more false alerts (more sensitivity).

### The Zoonotic Alert System

The methodology for generating the Z-Alert index integrates data management, a clustering technique, and multi-objective optimization. The proposed index-based alert system utilizes regional information to quantify the criticality of an event. Our clustering approach aims to identify broader spatiotemporal patterns of group of data points (a cluster). In this context, an event is defined as a cluster representing a phenomenon, based on occurrence patterns in both space and time. For each cluster formed, a numerical value is assigned, the Z-Alert index, reflecting its criticality, in which high values indicate high-risk regions. Fig 1 illustrates the main steps in the development of the zoonotic alert system.

**Figure 1.**
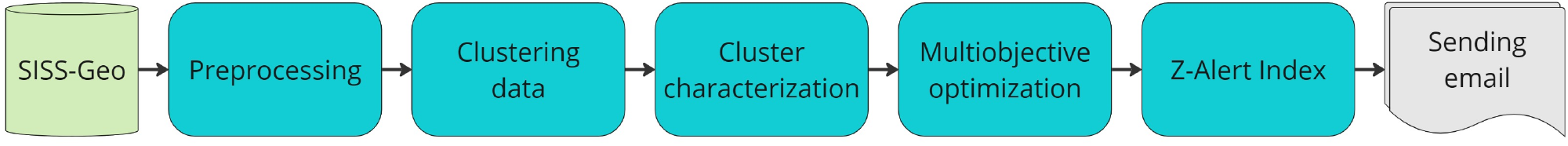
The zoonotic alert system. Main steps of the zoonotic alert system.

It is important to highlight that, although the national surveillance system recommends investigating a single dead or illness NHP, the alert index does not replace this approach—it complements it. While isolated cases are critical for early detection, clusters of NHP deaths often indicate that transmission is already underway. By clustering such occurrences and quantifying their attributes through the alert index, it becomes possible to prioritize areas where the risk is escalating, even when individual reports may seem sparse or dispersed.

Initially, data obtained from SISS-Geo are preprocessed according to the analyses of interest (section Data preprocessing). To extract regional information, a clustering technique will be applied, allowing similar instances to be grouped and organized into clusters (section Clustering data). After clustering, new attributes will be extracted from the records to characterize the clusters in greater detail (section Cluster characterization). In the optimization step (section Multiobjective optimization problem (MOP)), a multi-objective optimization process will be defined, parameterized by the cluster attributes, with the goal of obtaining an alert index that can effectively differentiate between critical and non-critical clusters. The optimization process will assign weights to each attribute, determining its relevance in calculating the Z-Alert index. Finally, the index is assigned to each cluster (section The Z-Alert Index), classifying them according to their level of criticality. Health sector managers will be able to define a threshold level for receiving alerts, which can be adjusted according to their needs and preferences. Clusters identified as critical will be highlighted and an automatic email will be sent to managers, informing them of the regions that require priority investigation.

A key advantage of the alert system proposed is its ability to provide a real-time risk signal, even in the absence of laboratory confirmation of circulating pathogens among animals. By leveraging spatio-temporal patterns extracted from the data, the index enables early identification of clusters with a high probability of being critical. This proactive approach offers an important decision-support tool for public health management, allowing preventive actions to be taken before zoonotic pathogens are confirmed, thus potentially reducing the time between emergence and response.

### Data preprocessing

The first step is the preprocessing of SISS-Geo’s data. The data collected are described in Table 1.

**Table 1:**
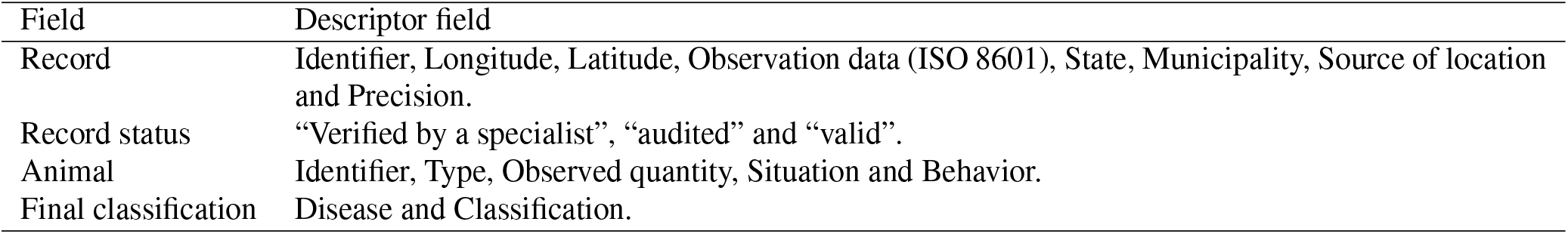
Data collected from SISS-Geo’s platform.

In the preprocessing stage, only records obtained via GPS or with locations explicitly provided by the user were selected, on top of having locations errors within 100m to ensure the spatial reliability of the data used for cluster formation. After data collection and filtering, each record was geocoded to its corresponding municipality using latitude and longitude coordinates, based on the official dataset provided by the Brazilian Institute of Geography and Statistics (IBGE) (available at https://geoftp.ibge.gov.br/organizacao_do_territorio/malhas_territoriais/malhas_municipais/municipio_2022/Brasil/BR/BR_Municipios_2022.zip). This information is necessary for validating the results presented in section Validation of the Z-Alert Index.

### Clustering data

Clustering techniques are designed to identify instances or groups of data that belong to the same phenomenon. The idea is to separate data with similar characteristics and assign them to a group or cluster. Given the dynamic nature of the phenomenon of interest, such as the occurrence of disease outbreaks—including YF—these events exhibit spatial and temporal movement patterns [9]. In this context, clustering enables the identification of groups of records that share these dynamic characteristics, supporting a more refined analysis of disease spread.

Several clustering algorithms can be found in the literature, the most traditional of which are available in the Scikitlearn open-source machine learning library. Fig 2 (sourced from https://scikit-learn.org/stable/modules/clustering.html#clustering) presents a comparative analysis of many of these methods applied to synthetic datasets with distinct cluster shapes. The patterns relevant to this study resemble those in the second and fourth rows of the figure, which depict curved clusters and clusters distributed along narrow bands. Among the evaluated methods, Spectral Clustering, DBSCAN, HDBSCAN, and OPTICS produced results most consistent with the expected cluster structure.

**Figure 2.**
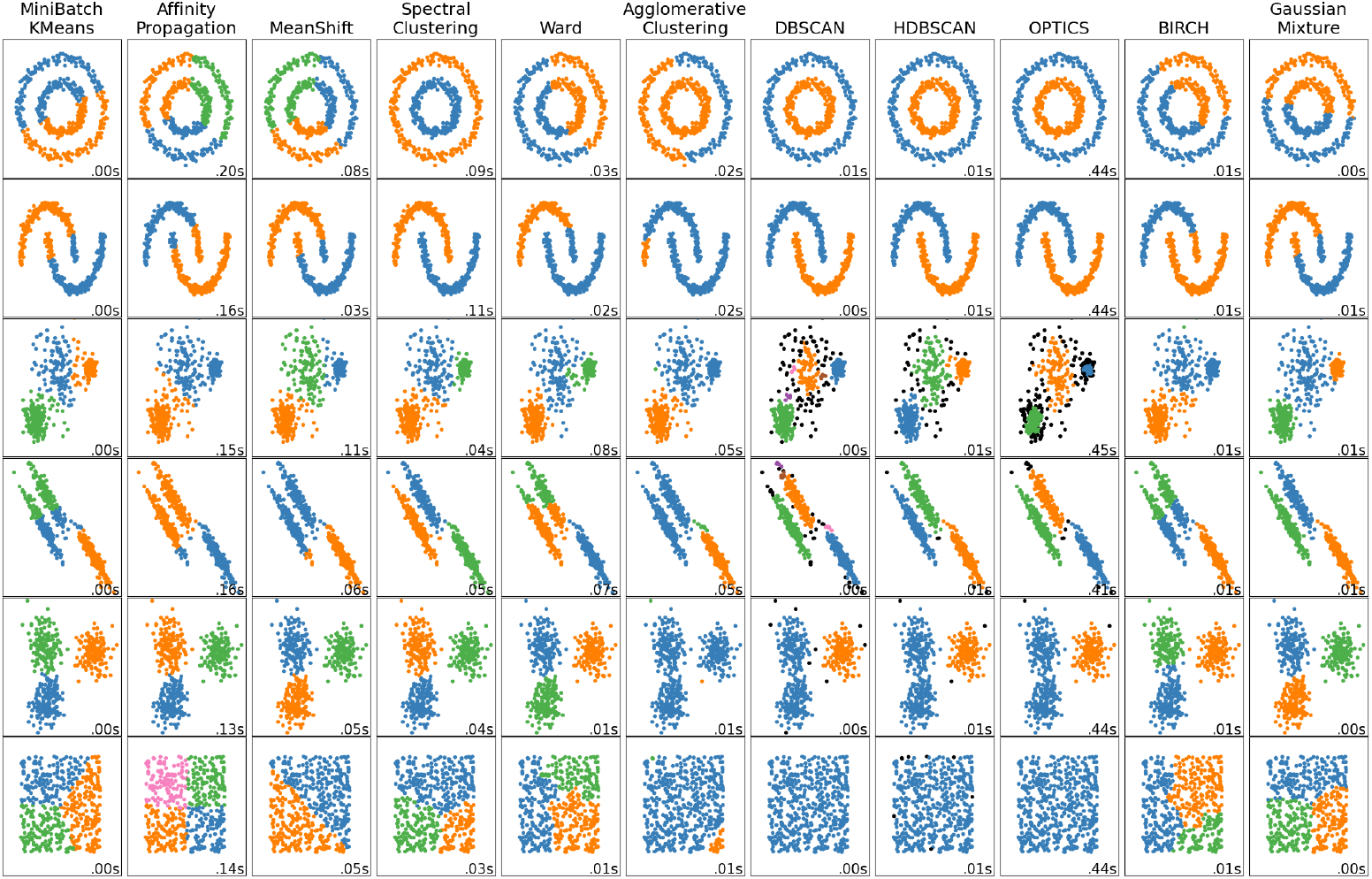
Clustering techniques. Comparative analyses of clusters techniques from Scikit-learn library.

From these, DBSCAN (Density-Based Spatial Clustering of Applications with Noise) [5, 21] was selected due to its ability to identify clusters of arbitrary shapes, its robustness to noise and outliers–data points that significantly deviate from the overall distribution–and its minimal parameter requirements. Additionally, unlike many clustering methods, DBSCAN does not require prior specification of the number of clusters, which is particularly advantageous in exploratory analyses where the underlying structure is unknown.

### The DBSCAN algorithm

The algorithm work by grouping points based on the density of neighbors within a specified radius (*eps*). It starts by selecting an unvisited point and checking whether it is a “core point”. If it is a core point, a new cluster is created, and all points within the *eps* radius are added to it. The process continues by expanding the cluster through neighboring points, checking their neighbors recursively until no further points satisfy the criteria defined by *eps* and the minimum number of points required to form a cluster. The algorithm then selects another unvisited point and repeats the process until all points have been visited. The parameters used in DBSCAN for the clustering step were set as follows:

*eps = 500*. Defines the neighborhood radius around a point, where two points are considered neighbors if the distance between them is less than or equal to *eps*. This value directly influences the size and density of the clusters.

*min samples = 1*. Represents the minimum number of points required within the *eps* radius for a point to be considered a core point and capable of expanding a cluster.

*metric = ‘precomputed’*. Specifies the distance metric used to calculate the proximity between points. In the experiments, instead of computing distances with a specific metric, the method directly receives a (square) matrix of precomputed distances between the points.

The choice of a high *eps* is justified by the need to generate clusters that follow the temporal and spatial definitions given by the precomputed distance matrix. Meanwhile, the minimum number of points required to form a cluster was set to 1 due to the nature of the SISS-Geo records, where isolated geographical points can be observed.

The distance matrix assigned to DBSCAN was generated by combining two matrices: (i) a spatial distance matrix *D_S_*, which considers the geodesic distance between points. The geodesic distance is calculated in a three-dimensional spherical space as the shortest path along the curved surface of the planet. This distance is computed based on the latitude and longitude of the points, and (ii) a temporal distance matrix *D_T_*, which indicates the time in days between records. Note that both *D_S_* and *D_T_* are square matrices, which is a prerequisite of the *metric* parameter.

The spatial and temporal constraints are considered when computing both matrices, with the temporal and spatial limits configured according to the experiments conducted (see section Results and Discussion). The matrices *D_S_* and *D_T_* are computed as follows. Let *D_S_*(*i, j*) be the geodesic distance between points *i* and *j*. If 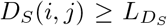, where 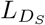 is the maximum spatial limit allowed for clustering points, a high value is assigned to indicate that these points cannot be grouped. Otherwise, *D_S_*(*i, j*) is assigned the actual distance between *i* and *j*. Similarly, let *D_T_* (*i, j*) be the temporal distance between instances *i* and *j*. If 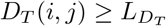, where 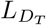 is the maximum temporal limit allowed for clustering, a high value is assigned to indicate that these instances cannot be grouped. Otherwise, *D_T_* (*i, j*) is assigned the time interval (in days) between *i* and *j*.

Since *D_S_* and *D_T_* have different scales (space versus time), both were normalized (values were scaled in the interval [0, 1]) to prevent large values from distorting the clustering process. The limits 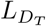 and 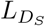 are defined by the user according to the experiments performed. Finally, the total distance matrix *D_total_* used as input for DBSCAN is computed as

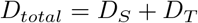

As a result of the clustering process, we obtain the clusters to which each record belongs, respecting the predefined temporal and spatial limits. It is important to highlight that the temporal and spatial limits are established for pairs of points, meaning that the generated cluster is not necessarily restricted to these limits.

### Cluster characterization

After the clustering step, new information is assigned to each cluster to support the construction of the Z-Alert index. The newly extracted attributes are presented in Table 2.

**Table 2:** New attributes extracted from clustering.

| Attribute | Description |
| --- | --- |
| $n\_death$ | Number of dead animals. |
| $n\_animal$ | Total number of animals involved; sum of live, sick and dead animals. |
| $ini\_date$ | Cluster start date, corresponding to the oldest record. |
| $end\_date$ | Cluster end date, corresponding to the most recent record. |
| $interval$ | Time interval (in days); difference between the dates of the earliest and most recent records. |
| $n\_rec$ | Number of records. |
| $freq\_rec$ | Frequency of records; number of records divided by $interval$. |
| $freq\_death$ | Frequency of dead animals; number of dead animals divided by $interval$. |
| $freq\_animal$ | Frequency of total animals; number of animals involved divided by $interval$. |
| $perc\_death$ | Percentage of dead animals; number of dead animals divided by $n\_animal$. |
| $extention$ | Maximum distance between all points in the cluster. |
| $n\_conf$ | Number of animals confirmed to have a disease of interest (e.g. YF). |
| $geocode$ | Cluster geocoding. |

The cluster geocode is determined based on the geocodes of the records it comprises. If a cluster contains records from different municipalities (i.e., with distinct geocodes), it is duplicated in the database to ensure that each entry maintains the correct association with its respective municipality. This information is only necessary to validate the results (see in section Validation of the Z-Alert Index).

### Multiobjective optimization problem (MOP)

Optimization refers to the study of problems in which the goal is to minimize or maximize a cost function, also known as the objective function, by selecting values for the function’s variables within a feasible set. In other words, among all possible solutions in the search space, we seek the one that best meets the desired criteria. In a single-objective optimization problem, only one function of interest is optimized. In this study, however, we aim to optimize multiple objective functions, mathematically formulated as [13]

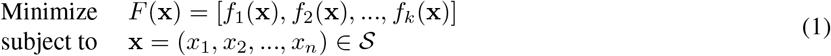

where *k*(≥2) is the number of objectives to be optimized, 𝒮 is called the feasible set, *F* is the vector of objective (function) values with *f_i_*: ℝ*^n^*→ ℝ, and **x** ∈ ℝ*^n^*is the vector of decision variables. If **x** ∈ 𝒮, then **x** is a feasible solution; otherwise, it is infeasible. The image of the feasible set, denoted by *Ƶ* (= *F* (𝒮)), is known as the feasible objective set. The elements of *Ƶ* are the objective functions with **z** = (*z*_1_*, z*_2_*, …, z_k_*)*^T^*, where *z_i_* = *f_i_*(**x**) for all *i* = 1*, …, k* are objective values. If an objective function *f_i_* is to be maximized, it is equivalent to minimize the function *f_i_*.

Due to the conflicting nature of the objective functions, which may also be in different units, it is not possible to find a single solution that is optimal for all objectives simultaneously. Instead, in MOP the aim is to identify a set of solutions, known as efficient or compromise solutions, that achieve the best possible trade-off among all objectives. This set of solutions is called the *Pareto-optimal solutions set*.

Assuming a problem in which all objectives should be minimized, a solution **x** ∈ 𝒮 dominates another solution **x***^′^* ∈ 𝒮 (written as **x** ≺ **x***^′^*), when

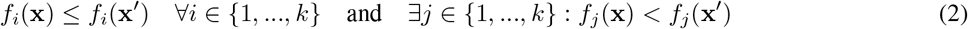

*i.e.*, the solution **x** is no worse than **x***^′^* in all objectives and better in at least one. Thus, all possible pairwise solutions can be compared in order to find which solutions are non-dominated. Finally, the set of non-dominated solutions is called the *Pareto-optimal set* and the corresponding image in the objective space defines the *Pareto-optimal Front*, representing the best collection of solutions found for the problem.

To obtain a representative set of solutions, in multi-objective optimization two orthogonal goals are desired: (i) find a set of solutions as close as possible to the Pareto-optimal front, and (ii) find a set of solutions as diverse as possible along the Pareto-optimal region [4]. Fig 3 illustrates a hypothetical Pareto front where objective *f*_1_ and *f*_2_ must be minimized. Converging to a set of solutions that are close to the true optimal and sparsely spaced in the Pareto-optimal region are the two main goals of MOPs.

**Figure 3.**
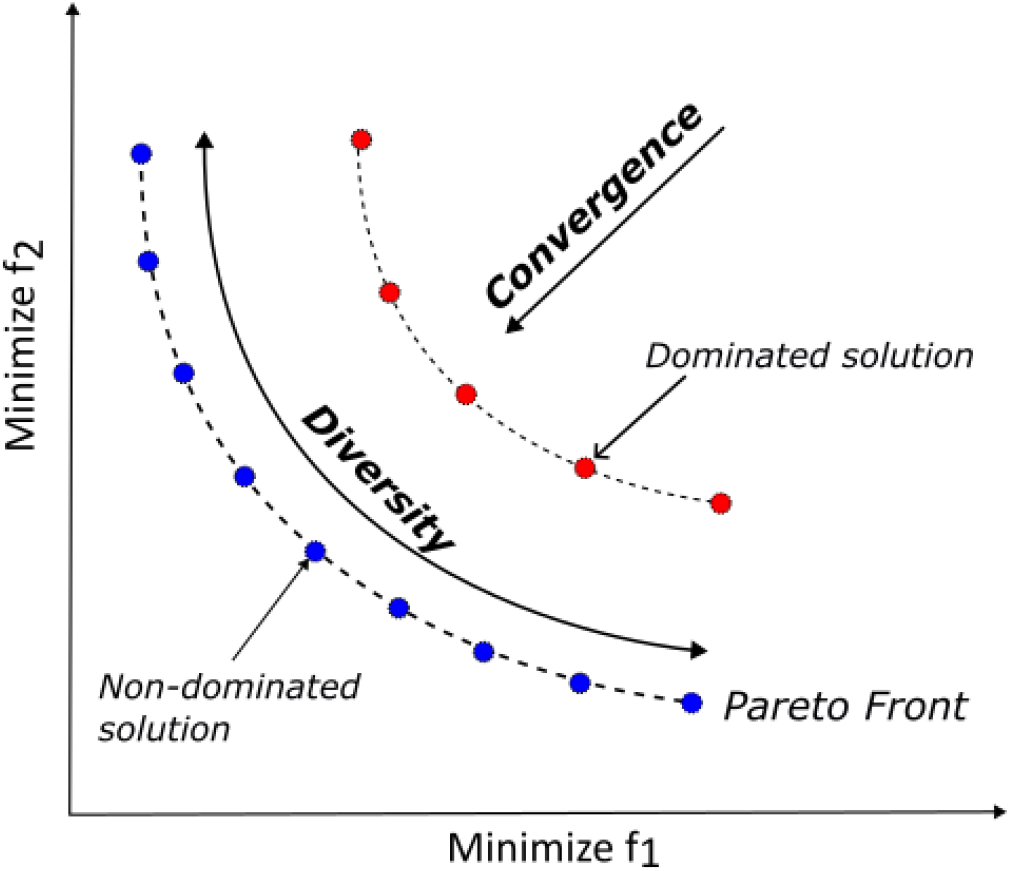
Two main goals of MOPs. Desired aspects of a Pareto front.

### Solution method

The simplest technique for solving MOPs involves aggregating the objectives using a weighted sum, transforming the MOP into a single-objective one. For each function *f_i_*, a weight *w_i_* is assigned such that 0 ≤ *w_i_* ≤ 1 for all *i* = 1*, …, k*, with 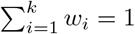. The transformed MOP is formulated as follows:

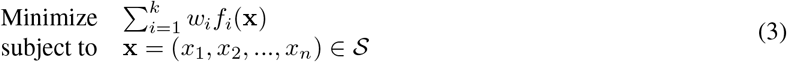

If all weight coefficients are positive or if the problem has a unique solution, then every solution found in (3) will be a Pareto-optimal solution [13]. The main advantage of this technique lies in its computational simplicity. However, obtaining the full set of Pareto-optimal solutions requires solving the optimization problem multiple times with different weight configurations. Once the Pareto set is obtained, the solutions are presented to the decision-maker, who selects the one that best aligns with their preferences. This approach was used to optimize the attribute weights that form the proposed Z-Alert index (in section The Z-Alert Index).

### Optimizing cluster attribute weights

The alert system currently implemented in SISS-Geo automatically generates an email whenever a dead or sick NHP is recorded. Since we aim to create an alert that anticipates the identification of phenomena of interest (e.g., a disease outbreak), only clusters containing more than one record were considered for the optimization process. This decision is motivated by the nature of the data registered in SISS-Geo, where most of the generated clusters—approximately 81% (see section Results and Discussion)—contain only a single record (*unit clusters*). If all of these clusters were included, the Z-Alert index would be biased toward unit clusters, which provide little regional information about the analyzed attributes compared to clusters with multiple records.

The goal of the Z-Alert index is to generate meaningful insights from data groups rather than individual records, enabling an accurate determination of a cluster’s criticality in relation to potential phenomena of interest. Thus, in the optimization phase, only clusters with more than one record were considered, and among these, only those containing confirmed cases of a disease of interest were selected. In the analyses conducted in section Analyzing the Pareto Front, we considered YF as the disease of interest. Clusters with confirmed cases were classified as *critical clusters*, while clusters without confirmed disease cases were classified as *non-critical clusters*. The formulated MOP aims to optimize two objectives simultaneously:

I. The mean of the critical clusters, which needs to be maximized, and
II. The variance of the critical clusters, which needs to be minimized.

The idea is to maximize a function that represents the combination of these two objectives. This ensures that the alert index effectively distinguishes critical clusters from non-critical ones while maintaining uniformity among the critical clusters. The objective function that we aim to maximize for optimizing the clusters’ attribute weights is

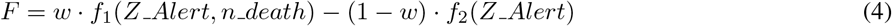

where *Z _Alert* (defined in the next section) are the indices of critical clusters, *f*_1_(*Z _Alert, n death*) represents the mean of the indices of critical clusters, weighted by the number of dead animals (*n death*), and *f*_2_(*Z _Alert*) represents the variance of the indices of critical clusters. The parameter *w* ≥ 0 controls the importance of maximizing *f*_1_, while (1 − *w*) controls the importance of minimizing *f*_2_, ensuring that *w* + (1 − *w*) = 1. Objectives *f*_1_ and *f*_2_ are computed as

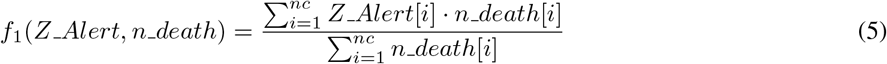

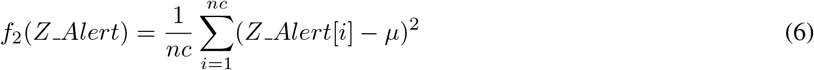

where *nc* is the total number of critical clusters and *µ* is the mean of *Z _Alert* of critical clusters. By maximizing the function (4), we encourage the attribute weights to adjust the index according to both combined objectives.

### The Z-Alert Index

For the construction of the alert index, let *M* be the attribute matrix of the clusters, composed of the columns

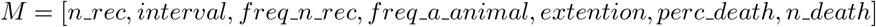

and let *p* ∈ ℝ*^m^*, where *m* is the number of variables, be the corresponding weight vector, which characterizes the importance of each attribute in computing the index. Due to the differences in scale among the variables, a data standardization process was applied to ensure that all variables contribute equally to the optimization step. The adopted standardization is based on the *Z-score* normalization, which transforms the data into a distribution with a mean zero and a standard deviation of one, calculated as

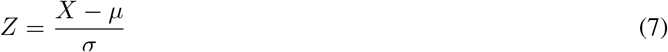

where *Z* is the normalized value, *X* is the original value of the variable, and *µ* and *σ* are the mean and the standard deviation of the variable, respectively.

Finally, for each cluster *i* = 1*, …, n*, where *n* is the total number of clusters, the *Z _Alert* index is calculated as

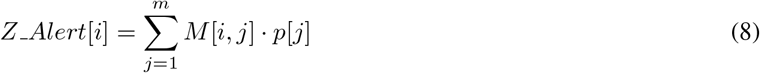

where *p*[*j*] is the weight associated to variable *j*, and *M* ∈ ℝ*^n×m^*is the matrix in which column *j* represents the value of the *j*-th variable and line *i* represents the *i*-th cluster.

At this stage, each cluster will receive a Z-Alert index, reflecting its level of criticality. Clusters with low index values are considered lower risk, while those with high index values indicate greater risk. However, the interpretation of this level of criticality may vary depending on the reality of each region, under the management of health departments. To accommodate this flexibility, the health manager can define a threshold level for receiving alerts, called *threshold level*, which can be adjusted according to their needs and preferences.

### The Threshold Level

The threshold level establishes a threshold for cluster criticality, directly influencing the number of alerts/notifications generated and ensuring that only the most relevant situations are reported to the manager. Considering that NHP mortality records depend on human analyses and may be subject to under-reporting, the alert index is designed to mitigate this by the aggregation of multiple attributes to provide a more robust signal. By calibrating the threshold appropriately, the system reduces the likelihood of overreacting to low-risk events while still capturing emerging patterns that may not be apparent from individual reports alone.

To define this threshold in an objective and adaptable manner, we use the distribution of the Z-Alert indices and perform the cutoff based on quartiles. Quartiles are statistical measures that divide an ordered set into four equal parts, each containing 25% of the observations. In the proposed scheme, quartiles are flexible, allowing the health manager to set the most appropriate threshold level for their reality. For instance, by choosing *threshold level* = 0.9 as the limit, only the top 10% most critical clusters will generate alerts, ensuring greater focus on high-risk regions. If a broader criterion is needed, the manager can opt for a lower quartile, increasing the number of received notifications. This approach provides greater flexibility and customization, allowing the system to adapt to different regional demands.

In the next section, the computational experiments will be presented, where different performance metrics were used to analyze the quality of the obtained solutions, evaluating the effectiveness of the Z-Alert index in distinguishing between critical and non-critical clusters and its impact on prioritizing notifications.

## Results and Discussion

In this section, we validated the results obtained by the Z-Alert Index using data from the Brazilian Ministry of Health (BMoH), which contains records of YF cases from 05 May 2014 to 31 December 2024 [15]. The initial date coincides with the first valid record from NHPs in SISS-Geo, while the final date reflects the ministry’s latest update on YF case occurrences in Brazil. The following criteria were considered for data collection:

Animal type: Marmoset and monkey.

Inclusion period: from 05 May 2014 to 31 December 2024.

Record status: “Verified by a specialist”, “audited” and “valid”.

Collection date: 04 April 2025.

As a result of this search, we obtained 6831 records. These data were filtered, considering only records obtained via GPS or explicitly provided by the user, with an accuracy of 100m, resulting in 4821 records for analysis. After data collection and filtering, each record was assigned the geocoding of the municipalities based on latitude and longitude information. This geolocation step was necessary only for validating the results, as the data provided by the BMoH reports YF cases at the municipal level.

Figs 4 and 5 present the clusters generated after the clustering step, using the limits 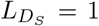 and 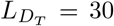. This process resulted in 3508 clusters, of which 2841 were unit clusters, representing approximately 81% of the total clusters. In the next stage, cluster characterization was performed, where new attributes were generated, as presented in section Cluster characterization.

**Figure 4.**
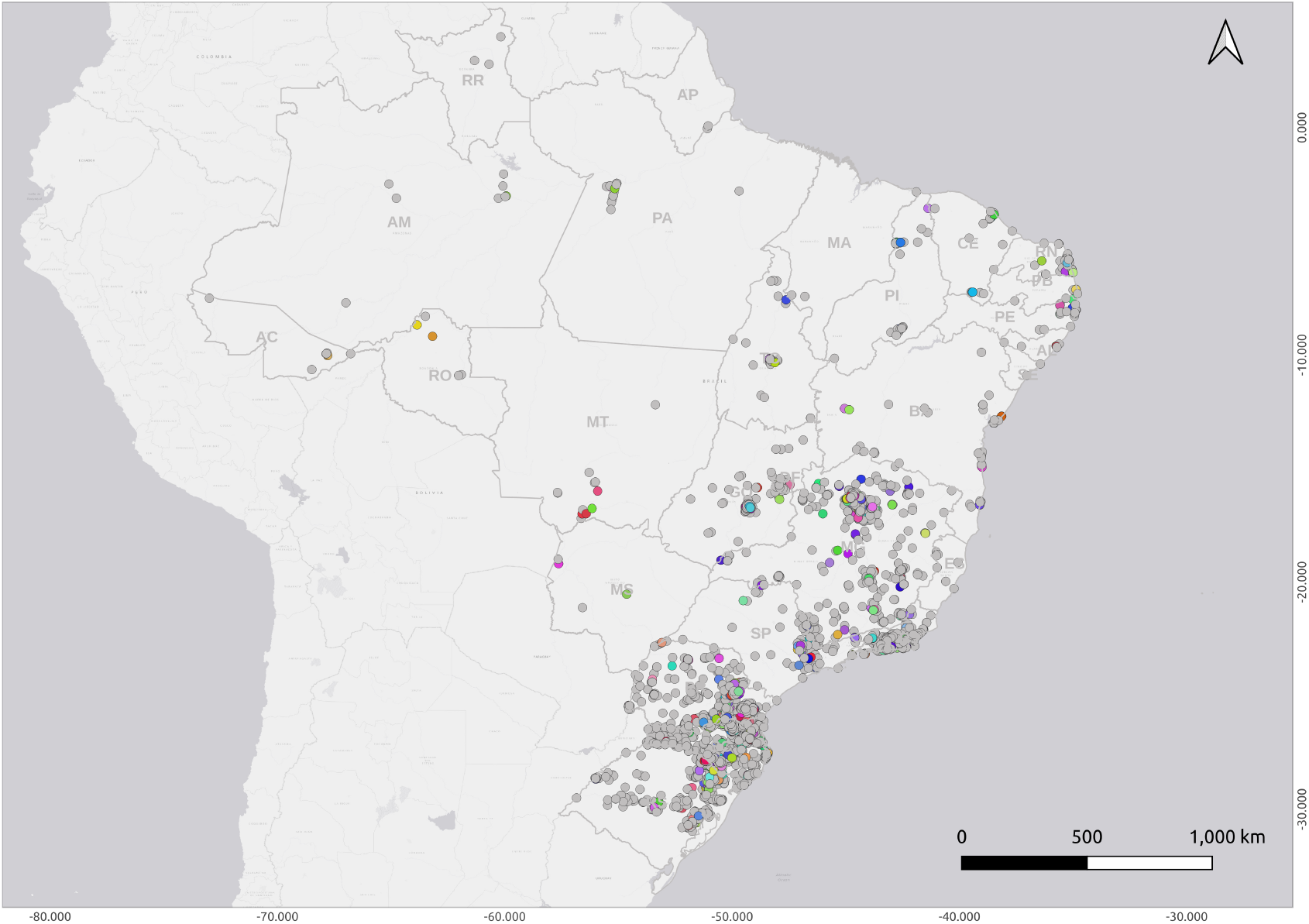
Data clustering. Illustration of all clusters obtained, where unit clusters are colored in gray circles.

**Figure 5.**
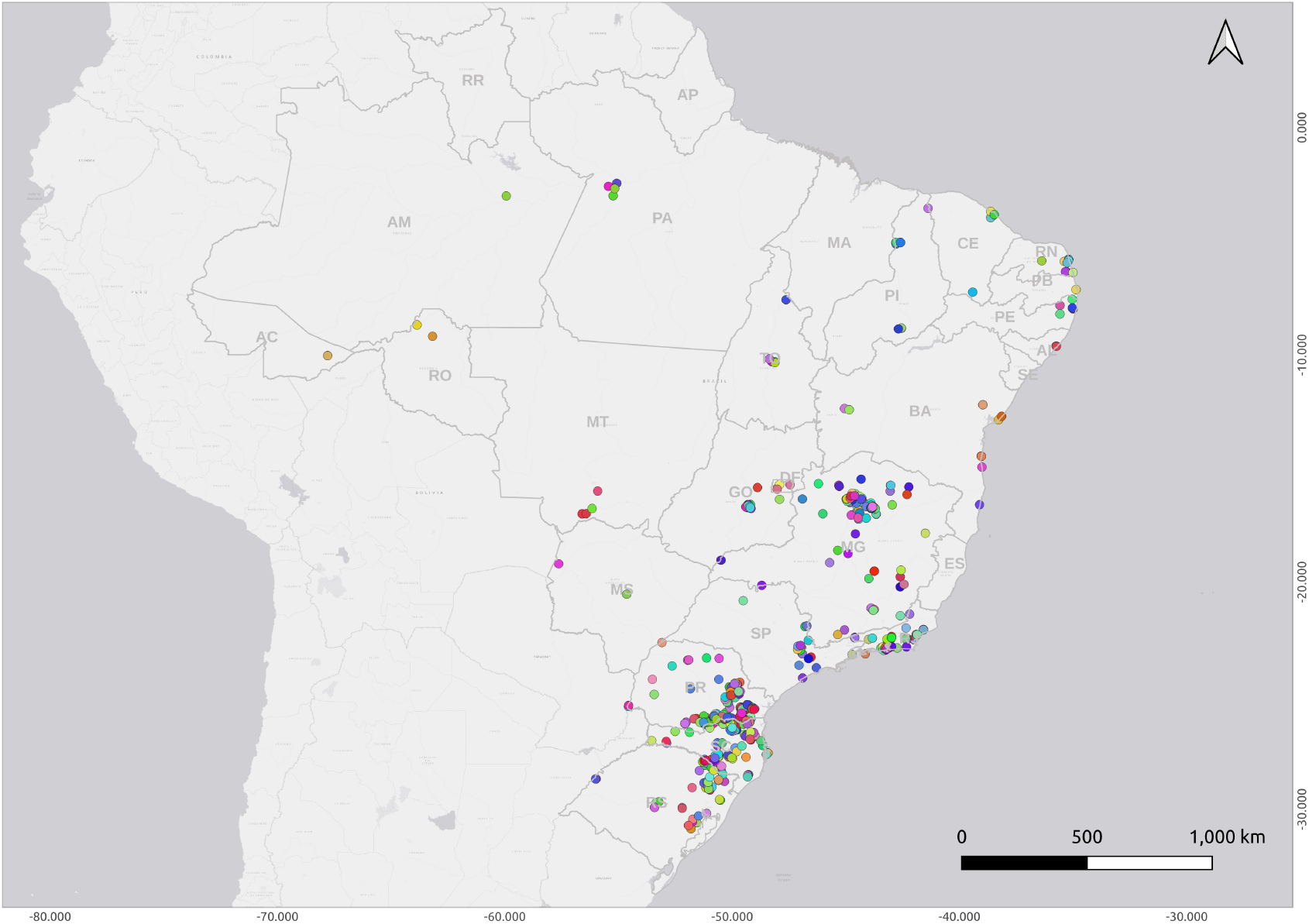
Data clustering. Emphasizing only non-unit clusters.

### Analyzing the Pareto Front

As described in section Optimizing cluster attribute weights, only clusters with more than one register were considered for the optimization process, and among these, only those containing confirmed YF records were included. Thus, for the attribute weights optimization process, 264 records grouped into 38 clusters were considered. To maximize the objective function *F* described in equation (4), the SLSQP (Sequential Least Squares Programming) method implemented in the SciPy library (https://scipy.org/) was used. The bounds on the variables are 0 ≤ *p_j_* ≤ 1 for all *j* = 1*, …, m*, constrained by 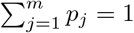. Different values of *w* were tested for generating the Pareto front. The parameter *w* controls the priority between the two optimization objectives:

- Higher values of *w* assign greater weight to the mean of critical clusters indices, emphasizing the maximization of alerts in high-risk regions.
- Lower values of *w* assign more importance to the variance of critical cluster indices, aiming to reduce value dispersion and ensure a more balanced distribution of alerts.

The tested values were

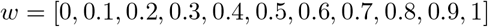

Additionally, we evaluated a scenario in which no optimization process was used for weight assignment. In this case, the weights were evenly distributed among all variables, assuming that each attribute has the same importance in constructing the Z-Alert Index. The SLSQP method was executed 11 times, once for each value of *w*, with *w* = 0 for the first experiment, *w* = 0.1 for the second, and so on. The non-optimized experiment was also considered, totaling 12 experiments. Each execution resulted in different values for the analyzed objectives. Fig 6 presents the obtained Pareto front, where objective *f*_1_ is maximized and *f*_2_ is minimized. The values from 1 to 12 indicate the solutions obtained, representing the experiments conducted.

**Figure 6.**
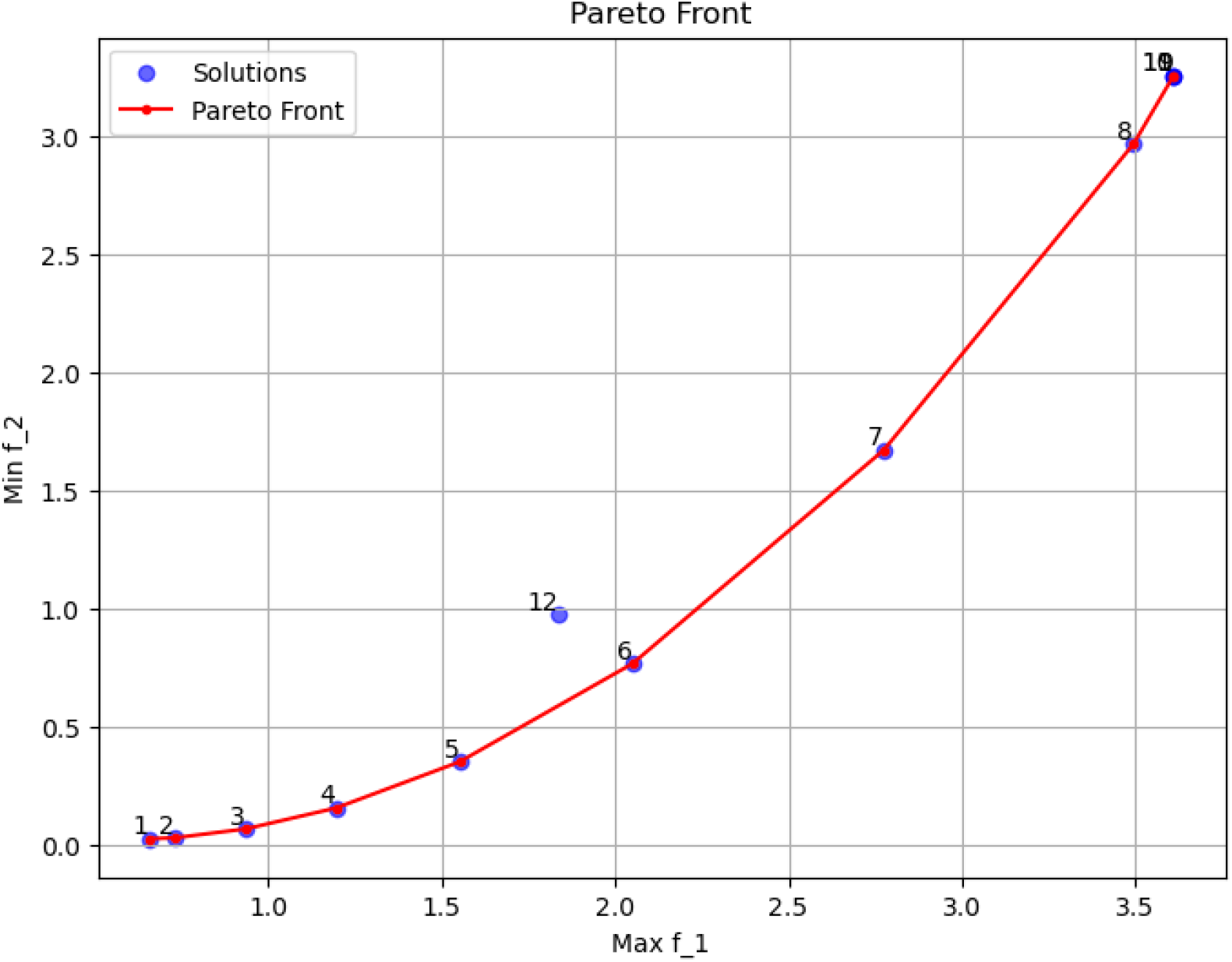
Pareto front. Pareto front obtained.

From these results, we can conclude the following points:

- The resulting front exhibits the expected behavior for a multiobjective optimization problem, where there is a trade-off between the two objectives [13]. As *f*_1_ improves (increases), *f*_2_ worsens (also increases). The red-connected points represent the Pareto front, meaning they are non-dominated solutions where no objective can be improved without compromising another. Note that uniformly distributing the values of *w* does not necessarily result in a uniformly distributed set of solutions on the Pareto front, as observed in the figure.
- The solution marked as 12 does not belong to the Pareto front, indicating that it is dominated by another solution. This means that assigning uniform weights to the attributes does not produce an efficient solution, reinforcing the importance of optimization in determining the weights.
- Solutions 9, 10, and 11, which correspond to *w* = 0.8, 0.9, 1, produced the same results. This suggests that the maximization of *f*_1_ reaches a limit, where further variations in *w* no longer affect model performance.
- The best solution depends on the application’s priority. If the goal is to only maximize *f*_1_, a point further to the right on the curve should be selected. If minimizing *f*_2_ is more important, the leftmost point is preferable. If the objective is to maintain a balanced index for critical clusters without excessively increasing variance, the optimal point should be near the middle of the curve, suggesting that experiment 6 is the best alternative, as it keeps *f*_1_ reasonably high while significantly improving *f*_2_.

### Analyzing the attributes’ weights

The attribute weights obtained in each solution are presented in Table 3. This table allows for a more detailed analysis of the relationship between attribute weights and objective values. From these results, we can highlight the following points:

**Table 3:** Assignment of attribute weights and objective values in each experiment.

| Sol. | $n\_rec$ | $interval$ | $freq\_rec$ | $freq\_animal$ | $extention$ | $perc\_death$ | $n\_death$ | $f_1$ | $f_2$ |
| --- | --- | --- | --- | --- | --- | --- | --- | --- | --- |
| 1 | 0.007 | 0.140 | 0.000 | 0.097 | 0.000 | 0.756 | 0.000 | 0.655 | 0.027 |
| 2 | 0.041 | 0.127 | 0.002 | 0.061 | 0.000 | 0.768 | 0.000 | 0.732 | 0.032 |
| 3 | 0.046 | 0.120 | 0.016 | 0.000 | 0.000 | 0.761 | 0.056 | 0.935 | 0.069 |
| 4 | 0.048 | 0.101 | 0.018 | 0.000 | 0.005 | 0.688 | 0.141 | 1.195 | 0.157 |
| 5 | 0.040 | 0.067 | 0.024 | 0.000 | 0.032 | 0.581 | 0.256 | 1.551 | 0.353 |
| 6 | 0.028 | 0.020 | 0.033 | 0.000 | 0.070 | 0.432 | 0.416 | 2.050 | 0.769 |
| 7 | 0.000 | 0.000 | 0.055 | 0.000 | 0.109 | 0.185 | 0.651 | 2.775 | 1.673 |
| 8 | 0.000 | 0.000 | 0.034 | 0.000 | 0.039 | 0.000 | 0.927 | 3.495 | 2.968 |
| 9 | 0.000 | 0.000 | 0.000 | 0.000 | 0.000 | 0.000 | 1.000 | 3.611 | 3.257 |
| 10 | 0.000 | 0.000 | 0.000 | 0.000 | 0.000 | 0.000 | 1.000 | 3.611 | 3.257 |
| 11 | 0.000 | 0.000 | 0.000 | 0.000 | 0.000 | 0.000 | 1.000 | 3.611 | 3.257 |
| 12 | 0.143 | 0.143 | 0.143 | 0.143 | 0.143 | 0.143 | 0.143 | 1.839 | 0.977 |

- Solution 12 (highlighted in the table) that applies uniform weights without optimization, presents a uniform distribution of weight values leading to a dominated solution in the objective space. This outcome highlights the importance of weight optimization, as it allows the alert index to better differentiate between clusters by emphasizing attributes that are more strongly associated with critical clusters. By assigning weights based on their contribution to relevant patterns, the optimized solution achieves improved performance in identifying high-risk areas.
- Solutions 9, 10, and 11 prioritize only the *n death* attribute, assigning 100% of the weight to it. These experiments achieve the best value for *f*_2_, indicating that *n death* is the most relevant attribute. However, since objective *f*_1_ worsens significantly, these solutions are not suitable if a trade-off must be maintained.
- Solution 7 and 8 prioritize the *n death* attribute but also introduce weights for *freq rec* and *extention*. The *perc death* attribute also shows relevance in experiment 7.
- Solutions 5 and 6 generate good trade-off solutions between *f*_1_ and *f*_2_. Both indicate that *perc deaths* plays a crucial role in balancing the objectives, with *n death* being the second most important attribute. Other attributes assigned weights in these experiments include *n rec, interval, freq rec*, and *extention*.
- Solutions 1 to 4 prioritize the minimization of *f*_2_, where *perc death* emerges as the most relevant attribute, followed by *interval, freq animal*, and *n rec*.

Observe that the attributes *n death* (number of dead animals) and *perc death* (percentage of deaths) capture complementary aspects of disease severity. While the absolute count provides a direct measure of mortality events, the latter highlights the lethality in proportion to the affected population, thereby enabling a more robust characterization of epidemiological risk. The presence of *perc death* seems to be crucial for determining the Z-Alert Index, as it appears to influence both objectives in a more balanced and controlled manner than *n death*. This outcome is expected, given the structure of the objective functions, which are formulated based on critical clusters. In these clusters, animals are confirmed to be infected with the YF virus and are therefore likely deceased. The other attributes show a certain degree of importance, appearing in a balanced way in most of the experiments.

### Validation of the Z-Alert Index

In this section, the Z-Alert Index is validated using data from the BMoH containing records of YF cases within a predetermined period. The goal is to verify whether the alerts generated by the index correspond to the cases recorded in the BMoH database. Validation will be carried out through the geocoding of municipalities and the analysis of the event occurrence period. To confirm an alert, we consider a 30-day window from the date of a confirmed case in the BMoH database. For instance, if the system generates an alert for a municipality on January 1, 2020, we check whether there is a confirmed YF case for that municipality in the BMoH database between that date and January 31, 2020, to confirm the alert in the corresponding cluster.

To assess the performance of the index, we use a confusion matrix [24]. The confusion matrix is a widely used tool to visualize the performance of classification algorithms. It organizes the results in a table that details the relationship between the model’s predictions and the actual observed values, as shown in Table 4.

**Table 4:** Confusion Matrix.

|  | Predicted Positive | Predicted Negative |
| --- | --- | --- |
| Actual Positive | True Positive (TP) | False Negative (FN) |
| Actual Negative | False Positive (FP) | True Negative (TN) |

In this study, the class defined as *actual* represents the data from the BMoH database containing confirmed YF cases. A municipality is classified as positive if a confirmed case of the disease is present, whether in humans or non-human primates. Thus, all records in the BMoH database correspond to positive cases. The class defined as *predicted* represents the alerts generated by the system. A cluster is classified as positive if it triggers an alert and negative otherwise. Based on this, the classification categories are defined as follows

- True Positive (TP): The system generates an alert, and the corresponding municipality has a confirmed YF case within the same period.
- False Negative (FN): The system does not generate an alert, but the corresponding municipality has a confirmed YF case within the same period.
- False Positive (FP): The system generates an alert, but the corresponding municipality does not have any confirmed YF cases within the same period. Notice that this is not a quite reliable metric due to probable under-reporting; in other words, the metric tends to be pessimistic.
- True Negative (TN): The system does not generate an alert, and the corresponding municipality also does not have any confirmed YF cases within the same period. Analogously, this metric may be impacted by under-reporting.

Based on this classification, various performance metrics were employed to evaluate the proposed alert system. The metrics used are described below:

- Precision: measures the accuracy of positive predictions, defined as: 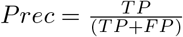
- Sensitivity: is the true positive rate, i.e., the proportion of correctly classified positive predictions, calculated as: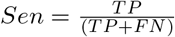
- F1-Score: corresponds to the harmonic mean between Precision and Sensitivity, expressed as: 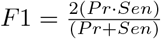
- Specificity: is the true negative rate, i.e., the proportion of correctly classified negative predictions, defined as: 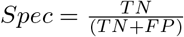
- Accuracy: represents the overall proportion of correct predictions, given by: 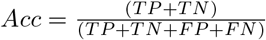
- Generalization: measures the model’s ability to correctly identify confirmed YF cases in relation to the total confirmed cases recorded in SISS-Geo, calculated as: 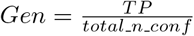

We evaluated the Z-Alert index for all clusters with more than one record whose municipalities and event periods match the information in the BMoH database. In this experiment, 492 clusters were assessed using different threshold levels, which define the criticality limit of the clusters (as detailed in section The Z-Alert Index). Table 5 presents the results of performance metrics for different threshold levels (0.8, 0.85, 0.9, 0.95, and 0.99) for the non-dominated solutions obtained in experiments 1, 6, and 11 (see section). These solutions were selected as the most representative based on the prioritization of objectives: solution 1 emphasizes minimizing *f*_2_, solution 6 aims to balance *f*_1_ and *f*_2_, and solution 11 focuses on maximizing *f*_1_. In table 5, TL indicates *threshold level*, Alert is the number of alerts, NonAL represents the number of non-alerts, and PAL corresponds to the proportion of alerts. The results in bold show the highest values.

**Table 5:** Performance metrics for the analysis of the Z-Alert Index for experiments 1, 6, and 11 presented in Fig 6.

| Sol. | TL | Alert | NonAL | PAL | TP | TN | FP | FN | Prec | Sen | F1 | Spec | Acc | Gen |
| --- | --- | --- | --- | --- | --- | --- | --- | --- | --- | --- | --- | --- | --- | --- |
| 1 | 0.8 | 349 | 143 | 0.71 | 188 | **124** | 161 | **19** | **0.54** | 0.91 | **0.68** | **0.44** | **0.63** | 5.22 |
| 6 | 0.8 | 364 | 128 | 0.74 | 193 | 114 | 171 | 14 | 0.53 | 0.93 | **0.68** | 0.40 | 0.62 | 5.36 |
| 11 | 0.8 | 387 | 105 | 0.79 | **203** | 101 | **184** | 4 | 0.52 | **0.98** | **0.68** | 0.35 | 0.62 | **5.64** |
| 1 | 0.85 | 341 | 151 | 0.69 | 182 | 126 | 159 | 25 | 0.53 | 0.88 | 0.66 | 0.44 | **0.63** | 5.06 |
| 6 | 0.85 | 355 | 137 | 0.72 | **189** | 119 | **166** | 18 | 0.53 | **0.91** | **0.67** | 0.42 | **0.63** | **5.25** |
| 11 | 0.85 | 297 | 195 | 0.60 | 159 | **147** | 138 | **48** | **0.54** | 0.77 | 0.63 | **0.52** | 0.62 | 4.42 |
| 1 | 0.9 | 230 | 262 | 0.47 | 131 | 186 | 99 | **76** | 0.57 | 0.63 | 0.60 | 0.65 | 0.64 | 3.64 |
| 6 | 0.9 | 229 | 263 | 0.47 | **136** | **192** | 93 | 71 | **0.59** | **0.66** | **0.62** | **0.67** | **0.67** | **3.78** |
| 11 | 0.9 | 232 | 260 | 0.47 | 132 | 185 | **100** | 75 | 0.57 | 0.64 | 0.60 | 0.65 | 0.64 | 3.67 |
| 1 | 0.95 | 132 | 360 | 0.27 | **78** | 231 | 54 | 129 | **0.59** | **0.38** | **0.46** | 0.81 | **0.63** | **2.17** |
| 6 | 0.95 | 135 | 357 | 0.27 | 77 | 227 | **58** | 130 | 0.57 | 0.37 | 0.45 | 0.80 | 0.62 | 2.14 |
| 11 | 0.95 | 116 | 376 | 0.24 | 66 | **235** | 50 | **141** | 0.57 | 0.32 | 0.41 | **0.82** | 0.61 | 1.83 |
| 1 | 0.99 | 24 | 468 | 0.05 | 12 | 273 | **12** | **195** | 0.50 | 0.06 | 0.10 | 0.96 | 0.58 | 0.33 |
| 6 | 0.99 | 27 | 465 | 0.05 | **18** | **276** | 9 | 189 | **0.67** | **0.09** | **0.15** | **0.97** | **0.60** | **0.50** |
| 11 | 0.99 | 26 | 466 | 0.05 | 17 | **276** | 9 | 190 | 0.65 | 0.08 | **0.15** | **0.97** | **0.60** | 0.47 |

From the results presented in Table 5, we can highlight the following points:

- Proportion of Alerts (PAL) and non-alerts: As the threshold level increases (from 0.8 to 0.99), the proportion of alerts decreases significantly, while the of non-alerts increases. This indicates that, with stricter thresholds, fewer clusters are classified as alerts. For example, at the 0.8 threshold, the PAL ranges from 0.71 to 0.79, while at the 0.99 threshold, PAL drops to 0.05.
- False Negatives (FN): Notice that the worst-case scenario occurs with the false negatives, as this indicates that the system failed to generate an alert for municipalities with confirmed YF cases. As the threshold level increases, false negatives also increase, because with higher thresholds, the model classifies fewer clusters as alerts.
- Precision (Prec): Precision remains nearly constant across all experiments and threshold levels, ranging from 0.52 to 0.65, indicating that, the proportion of correct alerts (with a confirmed YF case) remains relatively stable, with a slight improvement when threshold level increases.
- Sensitivity (Sen): Sensitivity drastically decreases as the threshold level increases. For instance, at the 0.8 threshold, sensitivity ranges from 0.91 to 0.98, while at the 0.99 threshold, it drops from 0.06 to 0.09. This shows that, with stricter thresholds, the model detects fewer alerts with confirmed YF cases, due to the overall reduction in the number of generated alerts.
- F1-Score (F1): The F1-Score, which balances precision and sensitivity, decreases as the threshold level increases, primarily due to a drop in sensitivity. At the 0.8 threshold, F1 is 0.68, while at the 0.99 threshold, it drops to 0.10 − 0.15. This indicates that, with stricter thresholds, the balance between precision and sensitivity decays.
- Specificity (Spec): Specificity increases with higher threshold levels, ranging from 0.44 (at the 0.8 threshold) to 0.97 (at the 0.99 threshold). This indicates that, with stricter thresholds, the model is better at correctly identifying non-alerts, likely due to the growing number of instances classified as non-alerts. It is important to highlight that specificity depends on false positives and true negatives, which both assume reliable knowledge of where YF did not occur. However, with under-reporting, some true cases go unrecorded. This makes specificity potentially overestimated or misleading.
- Accuracy (Acc): Accuracy is nearly constant across all experiments and threshold levels. This relatively constant values, despite large variations in sensitivity and specificity, may suggest that the model has limited discriminative power, that is, it struggles to effectively separate positive from negative cases. However, a more plausible explanation is that the accuracy metric is being disproportionately influenced by the majority class, probably due to a large number of true negatives, affected by under-reporting and the presence of undiagnosed or unconfirmed cases, an issue also emphasized in [6]. In imbalanced datasets, accuracy can remain misleadingly stable even when the model fails to correctly identify minority-class instances, in this context, true positive cases in disease detection scenarios.
- Generalization (Gen): Generalization decreases as the threshold level increases. This happens because, with stricter thresholds, the model classifies fewer clusters as alerts, resulting in fewer true positives. Here, it is interesting to note the trade-off between generalization and precision. Even though SISS-Geo database contains significantly fewer confirmed YF cases (185 records considered in the optimization process) compared to the BMoH database (3323 records), precision remains stable across all experiments and threshold levels, indicating the model’s robustness to variations in threshold levels.

Figs 7–9 illustrate the alert-generating clusters identified in solutions 1, 6 and 11 using a threshold level of 0.99. The color intensity reflects the alert index value, with darker shades indicating higher levels of risk associated with the clusters. The bottom right image in Fig 7 shows a cluster with records located along the border between two municipalities, Mafra and Itaiópolis in Santa Catarina state. In these situations, the alert is generated for both municipalities to ensure comprehensive coverage and response. In all figures, the effect of clustering the points according to the predefined spatial and temporal constraints (see in section The DBSCAN algorithm) is clearly visible, as the events are grouped into coherent regions that reflect both geographic proximity and timing, facilitating more accurate detection and analysis of potential outbreak patterns.

**Figure 7.**
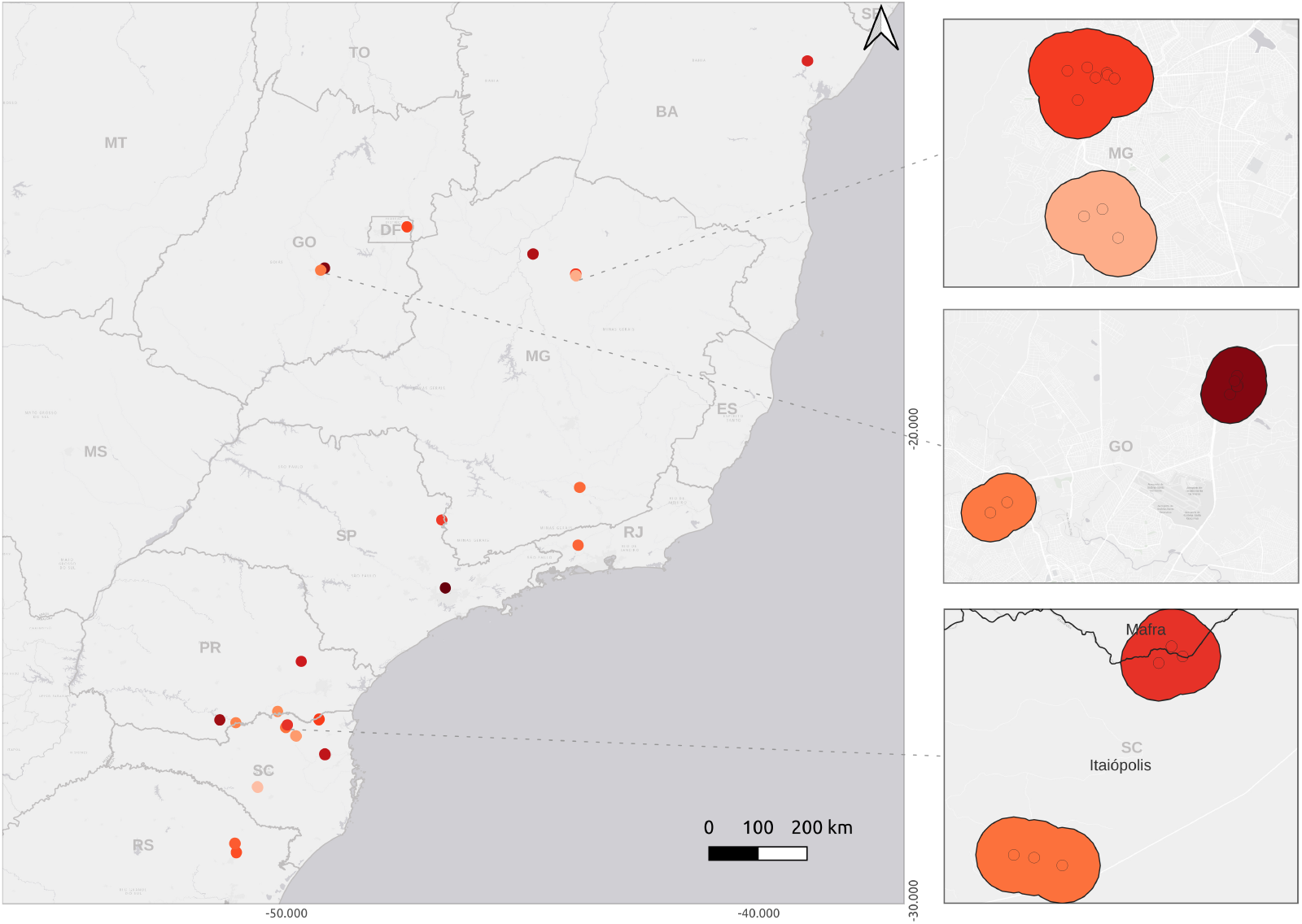
Alert-generating clusters. Result from solution 1 showing 24 alerts at threshold level 0.99.

**Figure 8.**
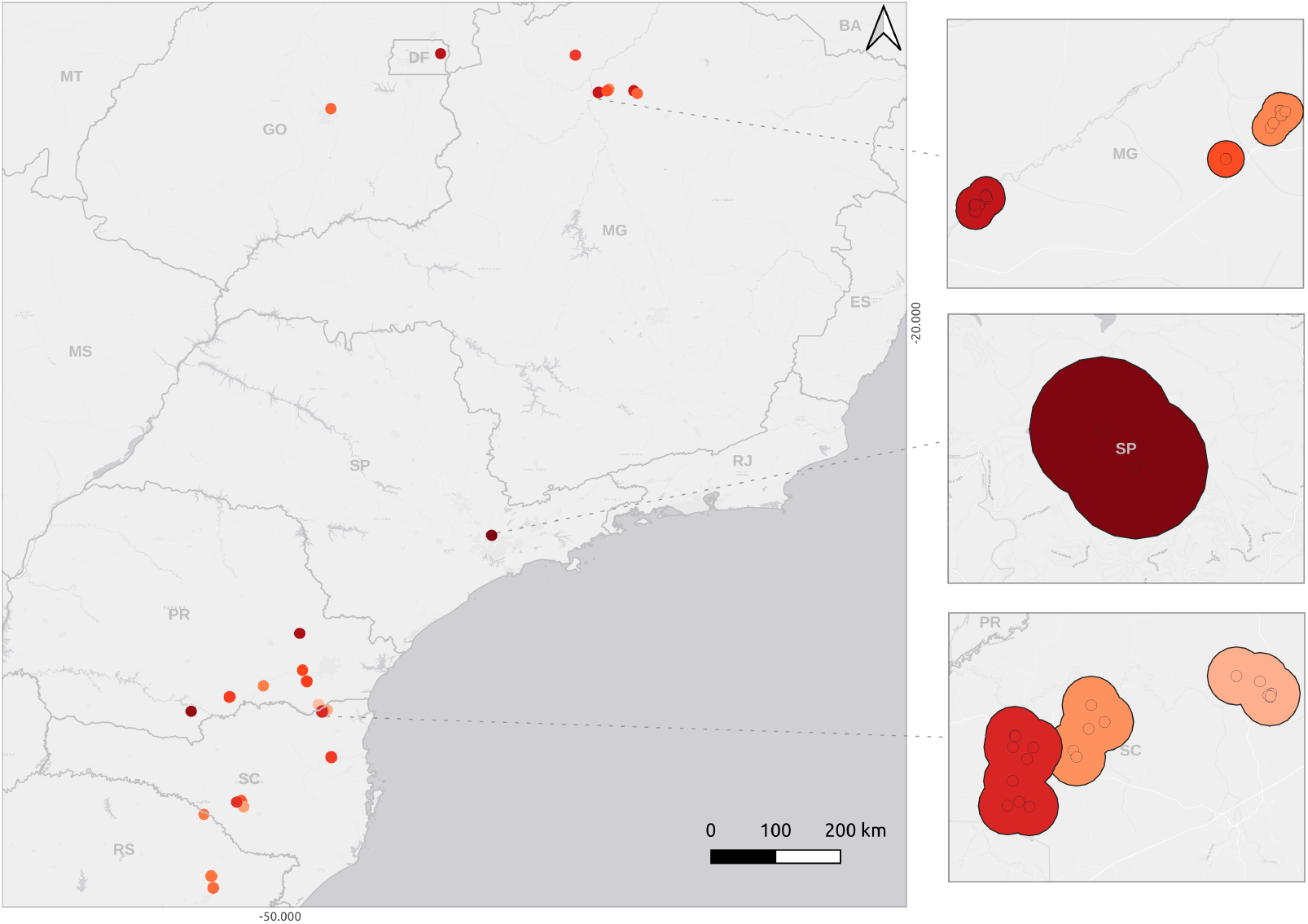
Alert-generating clusters. Result from solution 6 showing 27 alerts at threshold level 0.99.

**Figure 9.**
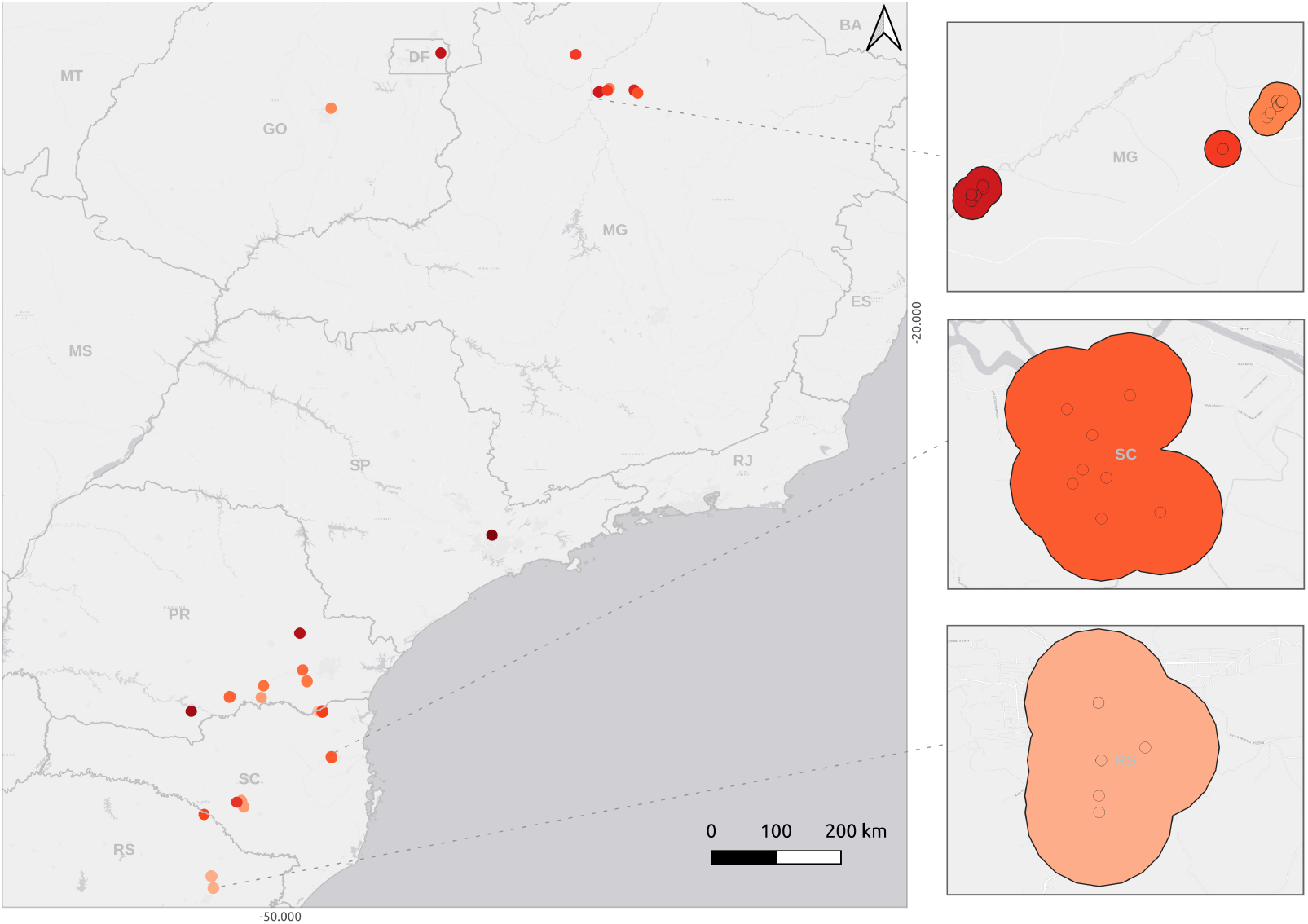
Alert-generating clusters. Result from solution 11 showing 26 alerts at threshold level 0.99.

By analyzing the results by solutions, a consistent trend is observed: as the threshold level increases, the number of alerts decreases, leading to higher specificity but lower sensitivity and F1-scores. Some other conclusions can be pointed out:

- Solution 1, which prioritizes uniformity across critical clusters, performance is characterized by the lowest precision (≈0.55 on average) across thresholds, compared with the other solutions. At TL = 0.8, this configuration achieves the heights precision, specificity and accuracy, but these values deteriorate as the threshold increases. This configuration is particularly suited for applications where false alerts carry a high cost, indicated by high specificity across the TL.
- Solution 6 seeks a balanced between the objectives and demonstrates stable and robust performance. At TL = 0.8, it attains good F1-score, with increased performance at TL=0.9 and 0.99. This configuration provides a good compromise between detection capability and reliability, making it appropriate when both avoiding false negative and capturing true cases are important.
- Solution 11, which aims to emphasize critical clusters, yields the highest sensitivity (0.98) at TL = 0.8 and the highest specificity at TL = 0.95 and TL = 0.99. This model prioritizes avoiding false positives even at the cost of missing true critical clusters. This setting is ideal in scenarios where missing false alerts has a greater cost than issuing true ones.

When comparing results by threshold level, lower TLs (0.8–0.85) result in a higher percentage of alerts (PAL = 0.71–0.79), higher sensitivity (0.91-0.98), and higher F1-scores (0.68), but also with more false positives. Conversely, higher TLs (0.95–0.99) drastically reduce the number of alerts (PAL = 0.27-0.05), improving specificity (up to 0.97 at TL=0.99) while severely compromising sensitivity (as low as 0.06) and F1-score (as low as 0.10). Higher thresholds are most appropriate when it is critical to avoid false alarms, even if most true cases are missed. Overall, Table 6 presents some recommended choices of threshold level (Rec. TL) based on defined objectives.

**Table 6:** Recommended choices of TL and solution based on defined objectives.

| Objective |  | Rec. TL | Solution | Justification |
| --- | --- | --- | --- | --- |
| Maximize detection of real cases |  | 0.8 | 11 | Highest sensitivity (0.98) and F1-score (0.68), minimal false negatives (FN = 4). |
| Best balance across metrics |  | 0.9 | 6 | More alerts, but all metrics balanced. |
| Minimize false negative |  | 0.85+ | 6 | On average was the lowest number of false negatives and high performance in all metrics on Rec. TL |
| Extremely conservative alerts |  | 0.99 | Any | Highest specificity (0.96-0.97), very few alerts (PAL = 0.05) and very low false alerts. |

From a practical perspective, defining the threshold level is essential, as it directly impacts the number of alerts that health managers will receive. Since this threshold can be adjusted to align with the specific needs and context of each region, it is important to ensure that only the most relevant situations are notified. Setting a higher threshold, resulting in fewer alerts, could be more appropriate in regions with high vaccination coverage and low human mobility, where the risk of disease spread is significantly reduced. If capturing as many alerts as possible is the priority, a lower threshold would be preferred, despite higher false positives may occur.

Based on the results, the solution that prioritizes the first objective (maximizing *f*_1_) is more effective at identifying real cases, although it may also produce a higher number of false positives. If a balance between detection and precision is required, aiming to reduce extreme variations, the solution seeking a balance between *f*_1_ and *f*_2_ stands out as the most appropriate alternative. On the other hand, if the priority is to minimize false alerts, even at the risk of missing some real cases, the solution that minimizes *f* 2 becomes the most suitable choice.

By observing the precision metric, it is possible to conclude that the model has a high ability to generate alerts for clusters with confirmed YF cases, even though this confirmation information is not present in the SISS-Geo database. The optimization process for setting the attribute weights was based solely on data from 38 clusters, which together account for 185 records with confirmed YF cases. This result indicates the model’s strong potential for generalization, successfully identifying critical clusters beyond those used during the optimization process. Notably, the alert index was also able to highlight high-risk clusters that did not contain confirmed cases, including some composed solely of live animals. In such cases, the alert system complements the current SISS-Geo rule-based approach, which would not have issued an alert under its fixed criteria.

It is also important to highlight the model’s limitations, which are directly related to the records provided by SISS-Geo. The effectiveness of the clustering process and the accuracy of the Z-Alert Index are highly dependent on the completeness, precision, and reliability of these records. Moreover, potential gaps in health surveillance, such as delays or failures in reporting confirmed YF cases or other diseases of interest, as well as the limited adoption of SISS-Geo in certain regions of the country, reduce the amount of available data. These issues hinder the model’s ability to detect meaningful patterns and to support the timely prioritization of preventive and investigative actions.

## Conclusion

The development of a multi-attribute zoonotic alert index (the Z-Alert Index) represents a significant advancement in epidemiological surveillance integrated within SISS-Geo. By adopting dynamic analyses based on clustering and optimization, the proposed system overcomes the limitations of the current model, enabling proactive identification of critical regions and efficient resource prioritization. Experiments with real-world data demonstrated the index’s ability to detect clusters associated with YF cases with high precision and adaptability to different scenarios. The flexibility to adjust parameters (such as spatial/temporal limits and threshold levels) ensures that the system can be customized according to regional specificities, diseases of interest and public health strategies.

Although the alert index showed strong performance in identifying clusters potentially associated with yellow fever virus circulation, its applicability to other zoonoses is constrained by data availability. While SISS-Geo also contains records related to rabies and avian influenza, these are considerably less frequent compared to yellow fever-related reports, which may limit the model’s sensitivity and generalizability when applied beyond the yellow fever context.

To address these limitations, future work will focus on expanding the model’s application to additional zoonoses— such as rabies and avian influenza—by both obtaining more data related to these diseases and integrating complementary environmental variables. Additionally, the optimization process will be continuously refined to reduce false negatives and enhance early warning capabilities across a broader range of epidemiological scenarios.

It is worth highlighting that regions with efficient monitoring and control strategies tend to attract more investments in technology innovation, and healthcare infrastructure. These benefits reinforce that investing in the identification and prioritization of critical regions is not only a public health concern but also a sound economic strategy. The proposed approach underlines the importance of collaboration between technology, data science, and healthcare management to address complex challenges at the interface of humans, animals, and the environment.

## Acknowledge

The authors thank the financial support provided by the Brazilian agencies Conselho Nacional de Desenvolvimento Científico e Tecnológico (CNPq) (grants 380656/2024-8 and 445493/2023-2).

